# Osteoporosis- and obesity-risk interrelationships: An epigenetic analysis of GWAS-derived SNPs at the developmental gene *TBX15*

**DOI:** 10.1101/766584

**Authors:** Xiao Zhang, Kenneth C. Ehrlich, Fangtang Yu, Xiaojun Hu, Hong-Wen Deng, Hui Shen, Melanie Ehrlich

## Abstract

A major challenge in translating findings from genome-wide association studies (GWAS) to biological mechanisms is pinpointing functional variants because only a very small percentage of variants associated with a given trait actually impact the trait. We used an extensive epigenetics, transcriptomics, and genetics analysis of the *TBX15/WARS2* neighborhood to prioritize this region’s best-candidate causal variants for the genetic risk of osteoporosis (estimated bone density, eBMD) and obesity (waist-hip ratio or waist circumference adjusted for body mass index). *TBX15* encodes a transcription factor that is important in bone development and adipose biology. Manual curation of 692 GWAS-derived variants gave eight strong candidates for causal SNPs that modulate *TBX15* transcription in subcutaneous adipose tissue (SAT) or osteoblasts, which highly and specifically express this gene. None of these SNPs were prioritized by Bayesian fine-mapping. The eight regulatory causal SNPs were in enhancer or promoter chromatin seen preferentially in SAT or osteoblasts at *TBX15* intron-1 or upstream. They overlap strongly predicted, allele-specific transcription factor binding sites. Our analysis suggests that these SNPs act independently of two missense SNPs in *TBX15*. Remarkably, five of the regulatory SNPs were associated with eBMD and obesity and had the same trait-increasing allele for both. We found that *WARS2* obesity-related SNPs can be ascribed to high linkage disequilibrium with *TBX15* intron-1 SNPs. Our findings from GWAS index, proxy, and imputed SNPs suggest that a few SNPs, including three in a 0.7-kb cluster, act as causal regulatory variants to fine-tune *TBX15* expression and, thereby, affect both obesity and osteoporosis risk.

## Introduction

Obesity is a public health problem affecting hundreds of millions of adults worldwide (1). It is a major risk factor for cardiovascular disease, chronic kidney disease, type 2 diabetes mellitus, and certain types of cancer, and thus poses a large and widespread health, social, and economic burden. The health consequences of obesity and, to a lesser extent, being overweight are determined not only by the total amount of body fat but also by where fat accumulates (1). Fat deposited around internal organs in the abdominal cavity (visceral adipose tissue, VAT) has a stronger association with metabolic and cardiovascular disease risk and mortality compared with hip or gluteal subcutaneous adipose tissue (SAT). VAT and SAT differ in preadipocyte replication and developmental origin, developmental gene expression, susceptibility to apoptosis and cellular senescence, vascularity, inflammatory cell infiltration, adipokine secretion and embryonic origin (2–5).

In the past 10 years, genome-wide association studies (GWAS) have identified more than 500 genetic loci associated with obesity-related traits. The studied traits are usually body mass index (BMI), which monitors overall obesity; waist circumference adjusted for BMI (WC_adjBMI_); and waist-hip ratio adjusted for BMI (WHR_adjBMI_), a proxy for body fat distribution (6–12). BMI-associated loci were enriched in genes expressed specifically in the central nervous system (8). In contrast, genes for loci associated with WHR_adjBMI_ showed enrichment in adipose tissue and in pathways for adipogenesis, angiogenesis, transcriptional regulation, and insulin resistance (9, 13).

The vast majority of risk variants (usually SNPs) from GWAS of obesity and other traits are located in non-exonic regions of the genome (8, 9, 12, 14). However, elucidating the causal SNP(s) and sometimes even the causal gene(s) within each obesity risk-related locus remains challenging (15). The GWAS-derived SNP exhibiting the smallest p-value for association with the studied trait at a given locus (index SNP) (16) is often linked to very many proxy SNPs solely due to linkage disequilibrium (LD) and not to biological significance. This plethora of proxy or imputed SNPs hinders the identification and functional characterization of the actual causal variants.

Recently, whole-genome profiles of epigenetics features, such as promoter-type or enhancer-type chromatin, have been used to help discriminate likely causal variants from bystander SNPs in high LD with them (15, 17, 18). GWAS SNPs, including those for obesity-related traits (9), are generally enriched in enhancer chromatin or promoter regions (14, 15, 19, 20). In GWAS of (mostly) individuals of European ancestry (EUR) (13, 21) or of East Asians (6, 11), SNPs linked to *TBX15* were identified as significantly associated with BMI, WHR_adjBMI_ or WC_adjBMI_ in mixed sex populations, or separately in women or men (Supplementary Material, Table S1). *TBX15* codes for a T-box transcription factor (TF) that may be involved in various aspects of adipose development and maintenance (22–24) and is known to be important in formation of the skeletal system (25, 26). SNPs in or near *TBX15* are significantly associated not only with obesity traits but also with estimated heel bone mineral density (eBMD), a useful indicator of hip and non-spine fractures that is used as a proxy for osteoporosis risk (27).

Our study of the relationships in GWAS of *TBX15* to both adipose and bone biology provides novel insights into the developmental relationships between two related tissue lineages converging on one gene. Both adipocytes and osteoblasts (ostb; progenitor cells for bone formation) can be formed from mesenchymal stem cells (MSC) depending on the induced TFs (28). There are associations between obesity and osteoporosis that are of practical importance although they are highly dependent in direction as well as extent on gender, age, and bone region susceptible to fracture (29, 30).

GWAS for obesity traits, although not for eBMD, have also identified several index SNPs in the gene body of *WARS2, TBX15*’s nearest-neighbor protein-coding gene*. WARS2* encodes a mitochondrial tryptophanyl tRNA synthetase that has been linked to mitochondrial-related neurodegeneration (31). One study suggested that increased *WARS2* expression is linked to decreasing visceral adiposity through effects on brown adipose tissue function (32). It is unclear which of the many obesity-associated SNPs in the *TBX15/WARS2* region or eBMD-detected SNPs at the *TBX15* locus are biologically relevant to obesity or osteoporosis.

Here we explore in depth the genetic, epigenetic, and transcriptomic evidence for a possible causal role of certain *TBX15-* or *WARS2-*associated genetic variants in obesity or osteoporosis risk. Our analysis prioritized eight SNPs from hundreds of GWAS-related SNPs in the *TBX15/WARS2* region as the most likely regulatory causal SNPs for obesity or osteoporosis risk. Importantly, half of these SNPs shared the same trait-increasing allele for both obesity and eBMD. These findings illustrate how manual curation of epigenetic and genetic data can help define candidates for causal GWAS-derived SNPs at a single gene locus of interest.

## Results

### The obesity- and eBMD-associated *TBX15* gene has highly tissue-specific expression unlike its neighbor, *WARS2*

*TBX15* is associated with WHR_adjBMI_, WC_adjBMI_ and BMI in many GWAS (Supplementary Material, Table S1). Of all the obesity-trait related genes, it had the highest preferential expression in SAT (but not VAT) relative to 34 types of non-adipose tissues as assessed by RNA-seq on hundreds of samples (GTEx; Fig. 1E and Supplementary Material, Table S2A and data not shown (20, 33)). The only tissues with higher expression than SAT of *TBX15* were skeletal muscle and tibial artery. The levels of expression of *TBX15* depend on the exact fat depot, as shown in mice (34). There are two well-annotated full-length (RefSeq) gene isoforms of *TBX15,* whose 5’ ends are separated by 1.7 kb (NM_001330677.1 and NM_152380.2). With the exceptions of testis and liver, the isoform with the proximal TSS (NM_001330677.1; Figs. 1A and 2A) predominated over the other isoform in all examined tissue samples (Supplementary Material, Table S2B). Interestingly, a third isoform (*ENST00000449873*) corresponds to just the 3’ end of canonical *TBX15* and is predicted to encode a truncated protein missing the DNA-binding domain. This isoform is the major one expressed in liver (Fig. 1A and C). Surprisingly, the RNA from this little-studied truncated isoform constitutes almost half of the *TBX15* RNA in SAT according to GTEx data.

**Figure 1.**
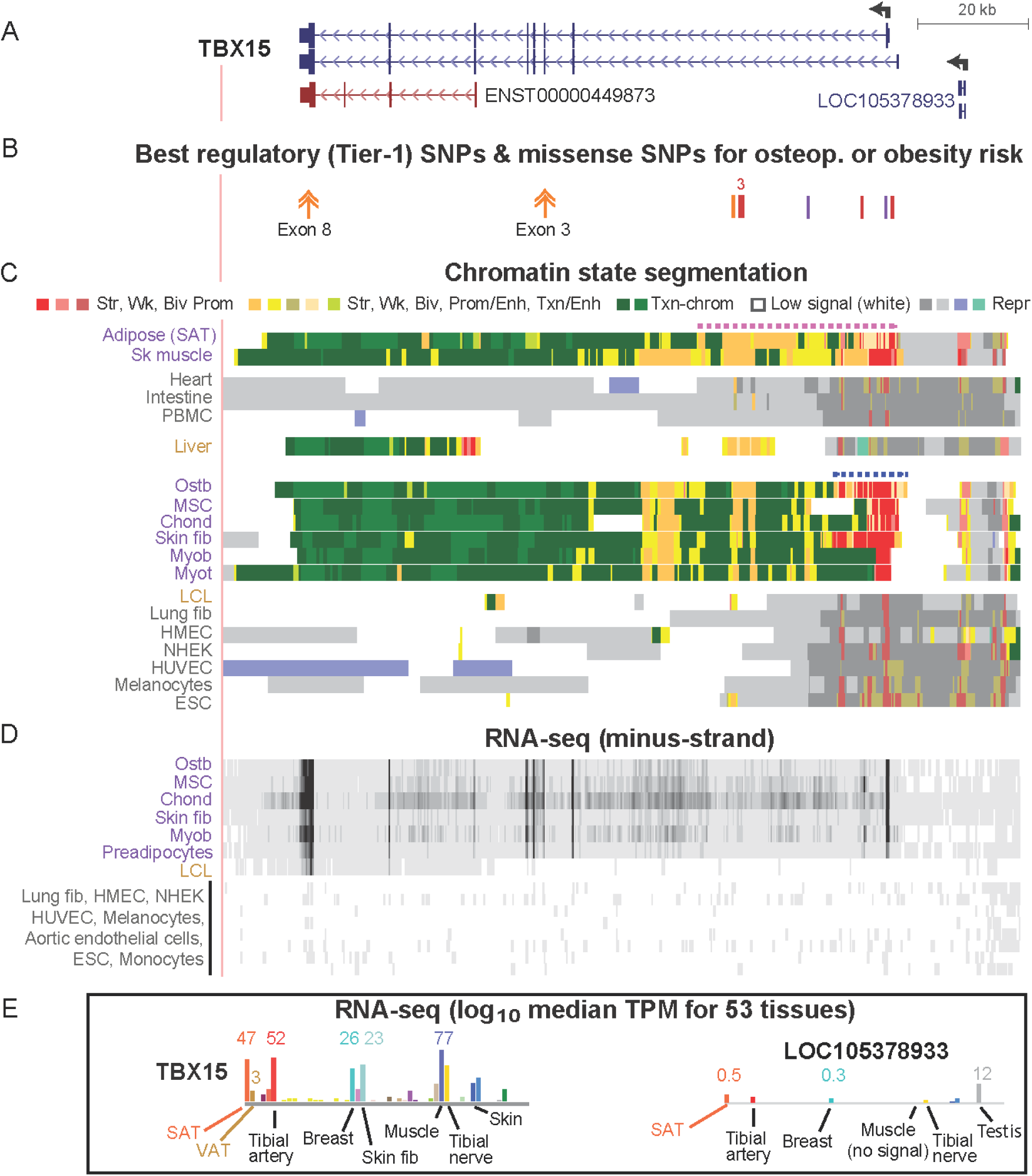
Obesity-trait or osteoporosis-related GWAS Tier-1 SNPs overlap tissue-specific regulatory chromatin in *TBX15*. (A) The two *TBX15* RefSeq gene isoforms, a truncated ENSEMBL isoform, and the upstream noncoding RNA gene (chr1:119,412,120-119,554,007, hg19); the broken arrow at *TBX15* shows the main TSS. (B) The location of two nonsynonymous coding SNPs (double arrows) and eight Tier-1 (best regulatory candidates) SNPs; “3,” a cluster of three Tier-1 SNPs. (C) Roadmap-derived chromatin state segmentation of strong (str), weak (wk), or bivalent (biv; poised) promoter or enhancer chromatin or repressed (repr) chromatin; SAT, subcutaneous adipose tissue; ostb, osteoblasts; sk muscle, skeletal muscle; PBMC, peripheral blood mononuclear cells; fib, fibroblasts; myob, myoblasts; myot, myotubes; LCL, lymphoblastoid cell line; HMEC, mammary epithelial cells; NHEK, embryonic kidney cells; HUVEC, umbilical vein endothelial cells; ESC, embryonic stem cells; dotted horizontal lines, super-enhancers. (D) ENCODE strand-specific total RNA-seq. (E) RNA-seq (log10) from GTEx; the median TPM (linear scale) is given for some of the samples; VAT, visceral adipose tissue. All tracks were visualized in the UCSC Genome Browser and are aligned in this figure and Figs. 2-5, and purple labels denote *TBX15*-expressing samples; gray, non-expressing samples; brown, samples expressing predominantly the truncated *TBX15* isoform.

*WARS2,* whose 3’ end is ∼45 kb upstream of the 5’ end of *TBX15,* has a broad expression profile among various tissues, a profile that is significantly different from that of *TBX15* (Table 1). *WARS2* is the nearest neighbor protein-encoding gene to *TBX15* and has been linked to obesity-trait risk in GWAS (6, 9) because of obesity-risk variants overlapping its gene body (Fig. 2C, triangles; Supplementary Material, Table S1).

**Figure 2.**
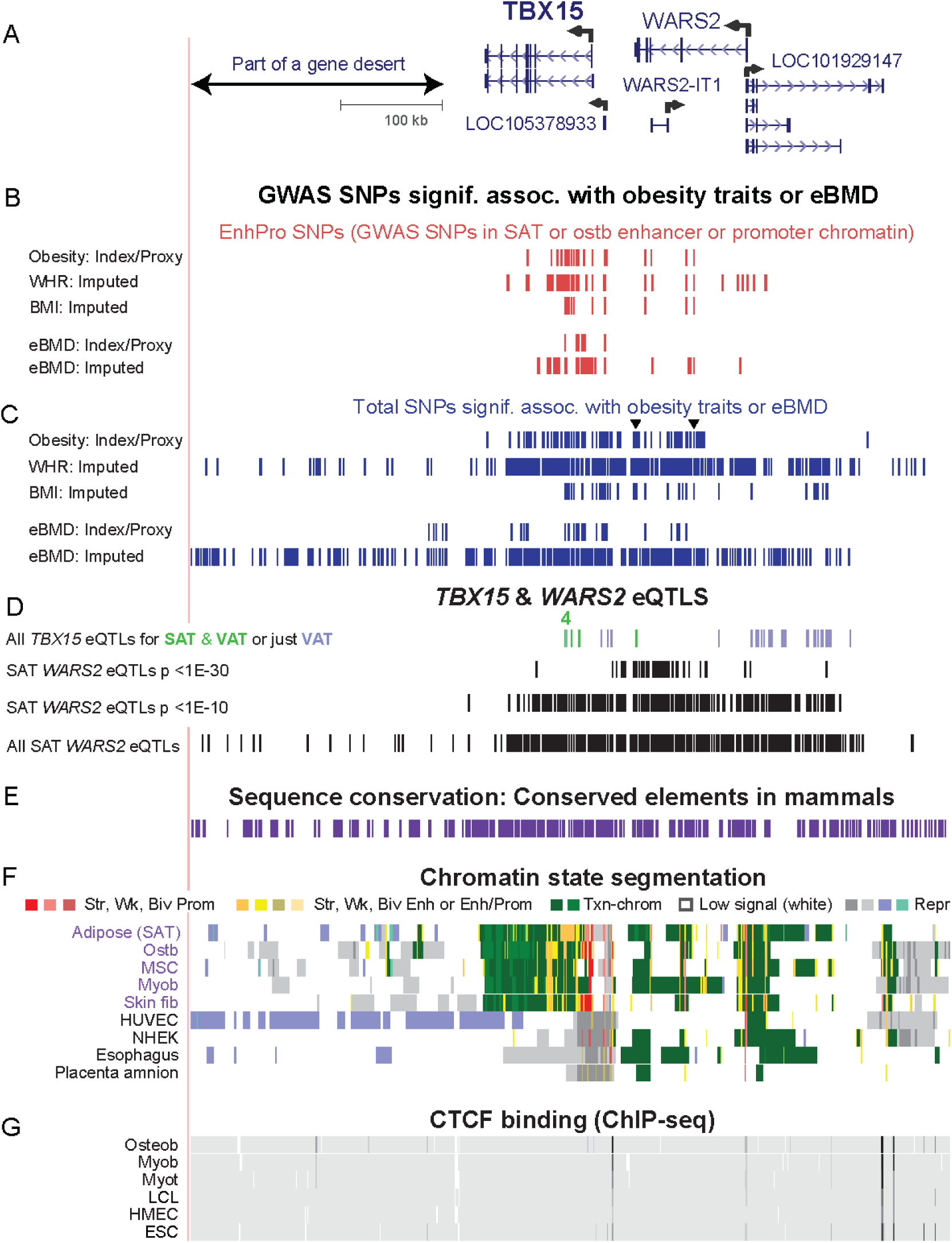
Many obesity-trait or eBMD risk-associated SNPs, EnhPro SNPs, and *cis*-eQTLs are found in the gene neighborhood of *TBX15.* (A) The gene neighborhood of *TBX15* (chr1:119,134,901-119,884,900). (B) Obesity-trait or eBMD GWAS SNPs are designated as EnhPro SNPs if they overlap enhancer or promoter chromatin preferentially in SAT or ostb. (C) All index SNPs, proxy SNPs (r^2^ ≥ 0.8, EUR) derived from them, and imputed SNPs in this region from obesity-trait or eBMD GWAS. (D) eQTLs for *TBX15* or *WARS2* in SAT or VAT. (E) Placental mammalian conserved elements (from phastCons track at the UCSC Genome Browser). (F) Chromatin state segmentation as in Fig. 1. (G) CTCF binding determined by Roadmap ChIP-seq. Panels B-D show custom tracks derived from Supplementary Material, Tables S4-S6.

**Table 1.**
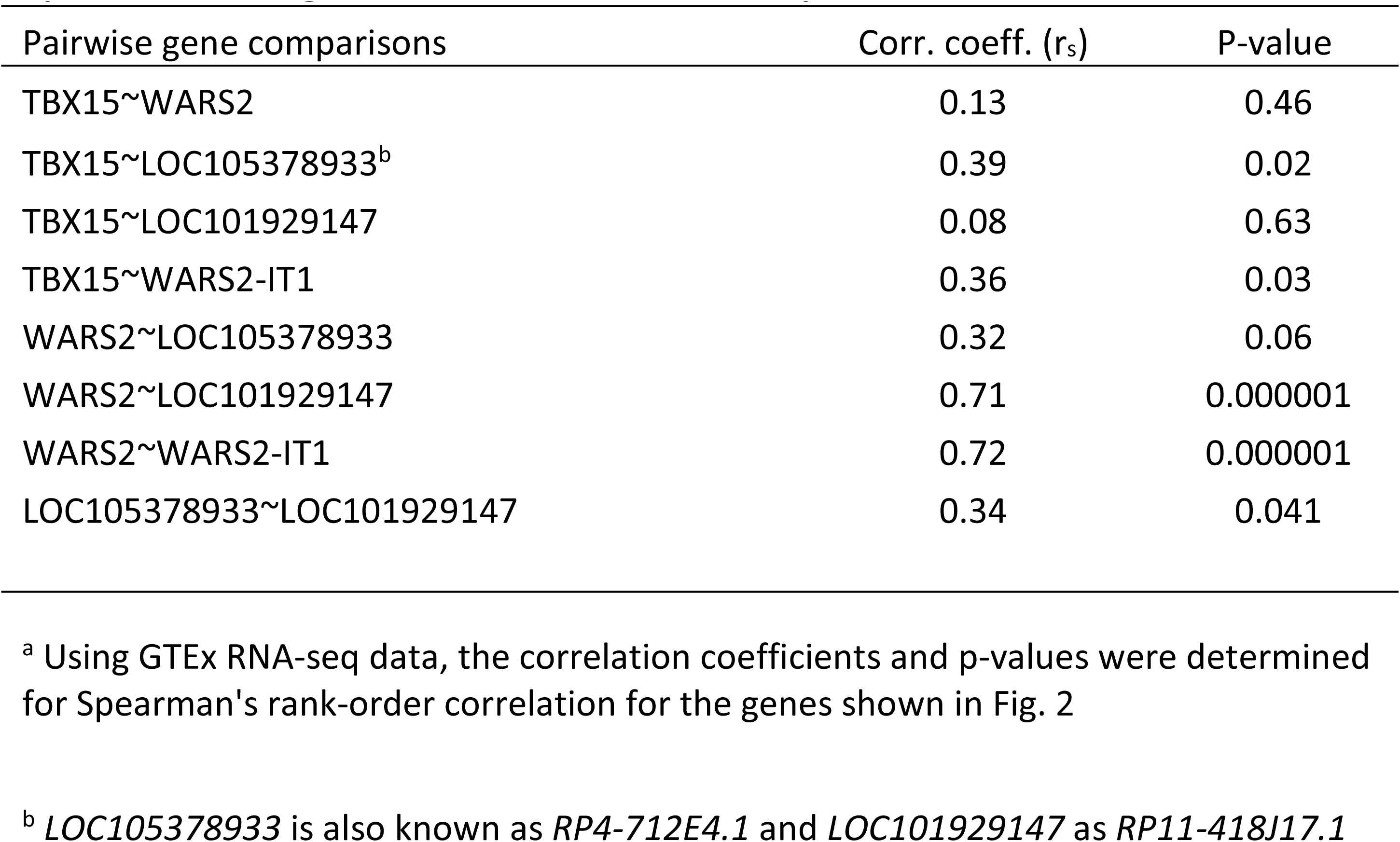
Expression of TBX15 is significantly associated with that of the immediately upstream ncRNA gene but not with the further upstream *WARS2*^a^

*TBX15* was significantly associated with eBMD, an indicator of osteoporosis risk in GWAS, as well as with obesity traits. Homozygous protein truncating mutations in *TBX15* are a cause of Cousin’s syndrome, a disease characterized by bone malformations and short stature (26), which is consistent with the involvement of this gene in osteoporosis. Steady-state RNA levels for *TBX15* in ostb are more than 200 times the median values for 11 heterologous cell strains (Fig. 1D) as seen in ENCODE RNA-seq data (35) (Supplementary Material, Table S3). MSC and chondrocytes (chond), both of which can be developmentally related to ostb (36), expressed this gene at similar high levels.

### High percentages of the index/proxy obesity- or BMD-related GWAS SNPs reside in regulatory chromatin in *TBX15* enhancer or promoter chromatin

To prioritize the best candidates for causal risk among the GWAS-derived SNPs in the vicinity of *TBX15* and *WARS2*, we used a multi-pronged approach. We assembled a set of significant (p < 5 x 10^-8^) obesity trait- and eBMD-related index SNPs at the *TBX15/WARS2* locus from many GWAS by querying the GWAS Catalog (37). The index SNPs were expanded to give proxy SNPs in high LD (r^2^ ≥ 0.8), and the sets of index and proxy SNPs were combined (index/proxy SNPs; Supplementary Material, Table S4). We separately collected sets of imputed SNPs (p < 6.6 x 10^-9^) at this locus obtained from summary statistics from one WHR_adjBMI_ meta-analysis of 694,649 individuals (38) and from an eBMD meta-analysis of 426,824 individuals (27) (Supplementary Material, Table S5). Because most of the GWAS used predominantly or only subjects of European ancestry, for all the linkage analyses described below we used only EUR populations. Among the index/proxy SNPs, 42% of the 109 obesity-trait SNPs (abbreviated as obesity SNPs) and 54% of the 37 eBMD SNPs overlapped the *TBX15* gene body while only 28% of the 427 WHR_adjBMI_ and 25% of the 443 eBMD imputed SNPs did (Fig. 2C). Most of the index/proxy obesity or eBMD SNPs were in the imputed WHR_adjBMI_, or eBMD SNP sets (72 and 76%, respectively). The more selective nature of the index/proxy set of SNPs might be a result of the GWAS Catalog extracting the most significant SNP from each independent locus to assign index SNPs (16).

We found that 38% (36) and 51% (19) of the obesity and eBMD index/proxy SNPs as well as 15% (65) and 10% (46) of the obesity and eBMD imputed SNPs in the *TBX15/WARS2* region overlapped enhancer/promoter chromatin preferentially in SAT or ostb, respectively (Fig. 2B and F; Supplementary Material, Tables S4 - S6). These variants will be referred to as EnhPro SNPs. Enhancer and promoter chromatin were defined by chromatin state segmentation in the Roadmap project as significantly enriched in histone H3 acetylation at lysine-27 (H3K27ac) and H3 monomethylation at lysine-4, (H3K4me1) or in H3K27ac and H3K4me3, respectively (14). The index/proxy or imputed EnhPro SNPs were found mostly in well-conserved intronic regions of *TBX15* (Fig. 2B and E).

The long intron 1 of *TBX15* was especially enriched in EnhPro SNPs. In SAT, they were mostly in the 36.5-kb super-enhancer overlapping this intron (Fig. 1C, dotted pink line). Most of the eBMD EnhPro SNPs in ostb were either in a smaller super-enhancer (Fig. 1C, dotted blue line) or in a downstream enhancer region in *TBX15* intron 1. Super-enhancers are unusually strong and long enhancers (> 5-kb) that contain clustered segments of chromatin enriched in H3K27ac and display the H3K4 methylation enrichment that is characteristic of enhancers or promoters (39).

### Obesity-related non-synonymous codon variants in *TBX15* exons 3 and 8

There are two obesity missense variants in *TBX15*, namely, rs10494217 (G/T, 0.84/0.16; H156N) in exon 3 and rs61730011 (A/C, 0.96/0.04; M566R) in exon 8 (Fig. 1B, double arrows) (13, 38, 40). The meta-analysis by Justice et al. (13) involving 476,546 individuals (88% EUR) found a highly significant association of both rs10494217 (p = 2 x 10^-14^) and rs61730011 (p = 3 x 10^-21^) with WHR_adjBMI_, and similar results were obtained for the former SNP in the study of Pullit et al. (38) (Supplementary Material, Table S5). Rs61730011 is in a codon for an amino acid in the large C-terminal portion of TBX15, which is of unknown function, and has no human paralogs (as determined by BLASTP, https://blast.ncbi.nlm.nih.gov/). In contrast, the amino acid associated with rs10494217 is in the T-box binding domain (41), and the SNP-overlapping codon and surrounding codons are highly conserved between *TBX18* and *TBX15*. Moreover, the corresponding amino acid sequences in these subregions of TBX15 display strong sequence conservation among mammals and vertebrates (Supplementary Material, Fig. S1), which suggests functionality of the corresponding protein subregion. Rs10494217 was predicted to have moderate impact on TBX15 protein function (40) even though an H → N substitution is a conservative one, unlike the rs61730011 M → R substitution. Mutations causing H → N substitutions in other proteins can be deleterious (42). Rs10494217, but not rs61730011, is also an eBMD-related SNP (Supplementary Material, Table S5). However, rs61730011 was indirectly implicated in bone biology by its significant association with adult height in a GWA study (43).

### Identification of eight best candidate regulatory SNPs (Tier-1 SNPs) for obesity-trait and/or eBMD risk

We stratified GWAS-derived *TBX15/WARS2* EnhPro SNPs to obtain a subset of best regulatory SNP candidates for obesity or osteoporosis risk (Tier-1 SNPs) by determining which of these SNPs that preferentially overlap enhancer or promoter chromatin in SAT or ostb also overlap DNaseI hypersensitive sites (DHS) in trait-relevant cell populations and predicted SNP-sensitive TF binding sites (TFBS). Because missense coding SNPs are especially good candidates for risk-modifying alleles, we first eliminated from further consideration any EnhPro SNPs that were in LD with r^2^ > 0.2 to either of the two above-mentioned missense obesity SNPs (Supplementary Material, Fig. S2). Next, for an EnhPro SNP to be considered a Tier-1 SNP, it had to overlap a DHS in SAT, adipocytes, or ostb. We included the frequently observed overlap with only a small DHS or the shoulder of a larger DHS rather than requiring the SNP to be embedded in the central region of a moderate-to-large DHS peak. In addition, we required that Tier-1 SNPs were located at a predicted TFBS that should exhibit allele-specific binding to a TF expressed in ostb or SAT. Allele-specific TFBS were identified using four TFBS prediction programs, JASPAR, TRANSFAC, HaploReg, and SNP2TFBS (44–47) (Supplementary Material, Table S7) and stringent manual curation (see Methods). We also looked for evidence of *in vivo* binding from genome-wide TF chromatin immunoprecipitation (ChiP-seq; Factorbook and UniBind (48, 49)) but those databases were informative for only one of the identified Tier-1 SNPs (rs12742627, Table 2) probably because of the lack of ostb and SAT samples in the database and the tissue-specificity of *TBX15 cis*-regulatory elements.

**Table 2.** Eight best candidates for transcription-regulatory obesity- or eBMD-causal SNPs (Tier-1 SNPs) in the *TBX15* neighborhood^a^

| Tier-1 SNP for obesity (ob) or BMD assocn.<sup>b</sup> | Index/proxy or imputed SNP | P-value for imputed assocn-with WHR or eBMD | SNP's region within *TBX15* | SNP's distance to the main *TBX15* TSS (kb) | Alleles (Ref/Alt) | Minor allele frequency | Trait-increasing allele (frequency)<sup>c</sup> | Chromatin state at the SNP in SAT and osteoblasts<sup>d</sup> | Predicted TF binding<sup>e</sup> | | Type of DHS at the SNP in SAT/adipocytes and ostb<sup>d</sup> | LD thinned<sup>f</sup> SNP group ($r^2 > 0.2$) for obesity | LD thinned<sup>f</sup> SNP group ($r^2 > 0.2$) for BMD |
| --- | --- | --- | --- | --- | --- | --- | --- | --- | --- | --- | --- | --- | --- |
|  |  |  |  |  |  |  |  |  | Ref allele | Alt allele |  |  |  |
| rs10923711 | Imputed (ob) | 6.5E-9 (WHR) | Intron 1 | 27.6 | T/C | 0.20 | Alt (0.20) | Str enh | ETS1, ETV5 ELK1 | None | Medium | IV | NA |
| rs984222 | Index/proxy (ob) Imputed (ob & BMD) | 3.8E-83 (WHR)<br>7.1E-18 (BMD) | Intron 1 | 26.6 | C/G | 0.39 | Alt (0.61) | Str enh | MZF1, ZNF281 | None | Small | I | I |
| rs9659323 | Index/proxy & imputed (ob & BMD) | 1.4E-48 (WHR)<br>1.5E-33 (BMD) | Intron 1 | 26.1 | A/G | 0.41 | Alt (0.41) | Str enh | AP-1 family, MAF/NFE2 | None | Small | I | I |
| rs1409157 | Index/proxy (ob) Imputed (ob & BMD) | 3.7E-71 (WHR)<br>1.2E-17 (BMD) | Intron 1 | 26.0 | G/T | 0.39 | Alt (0.61) | Str enh | GATA2 (SAT), GATA6 (ostb) | None | Shoulder | I | I |
| rs61806235 | Imputed (BMD) | 5.6E-19 (BMD) | Intron 1 | 14.0 | A/G | 0.10 | Alt (0.10) | Wk enh | None | AHR | Large | NA | II |
| rs12742627 | Imputed (ob & BMD) | 4.2E-11 (WHR)<br>2.4E-10 (BMD) | Intron 1 | 4.7 | G/A | 0.06 | Ref (0.94) | Str pro | TFDP1, E2F1, E2F4, E2F6 (UniBind) | None | Medium | V | III |
| rs961470 | Imputed (BMD) | 1.0E-10 (BMD) | Intron 1 | 0.3 | A/G | 0.06 | Ref (0.94) | Str pro &/or enh | None | NFIC, SMAD3 | Medium | NA | III |
| rs1106529 | Index/proxy (ob) Imputed (ob & BMD) | 1.2E-80 (WHR)<br>3.4E-17 (BMD) | TSS -1 kb | -1.0 | G/A | 0.27 | Alt (0.73) | Str pro | None | FOXO1, FOXC2 SOX4 | Small | I | I |

<sup>a</sup>Regulatory SNPs in the neighborhood of *TBX15* were imputed SNPs for WHR<sub>adjBMI</sub> from Pulit et al. (38) or for eBMD from Morris et al. (27) GWAS or were index plus proxy SNPs from BMI, WHR<sub>adjBMI</sub> and WC<sub>adjBMI</sub> or eBMD (see Supplementary Materials, Table S1). Not included in this table are two exonic missense variants with the respective trait-increasing alleles rs10494217-T (obesity and eBMD GWAS; Alt allele) and rs61730011-A (obesity GWAS; Ref allele); all p-value and LD data are for the EUR population (76); NA, not applicable
<sup>b</sup>Tier-1 SNPs are associated with allele-specific predictions of TF binding sites and overlap enhancer (enh) or promoter (pro) chromatin and DNaseI hypersensitive sites (DHS) preferentially in subcutaneous adipose tissue (SAT)/adipocytes or osteoblasts (ostb); rs984222 and rs1409157 are in perfect LD, and rs12742627 and rs961470 are in almost perfect LD ( $r^2 = 0.98$ )
<sup>c</sup>The eBMD increasing allele and the obesity-trait increasing allele was the same for both types of traits for alleles associated with obesity as well as eBMD
<sup>d</sup>Chromatin state and size of DHS from Roadmap for SAT/adipocytes or ostb; str, strong; wk, weak
<sup>e</sup>Predictions of allele-specific TFBS by hand curation of data from four programs (see Supplemental Material, Table S7) and assessment of TF expression in the target tissue
<sup>f</sup>LD thinning by LDLink SNPclip program (52); see Fig 6

Only eight of 93 obesity or eBMD EnhPro SNPs (Supplementary Material, Table S6) satisfied all the above criteria for designation as Tier-1 SNPs (Table 2, Figs. 3-5). Five were both obesity and eBMD SNPs, one was obesity only, and two were eBMD only. Most of the *TBX15* Tier-1 SNPs associated with obesity or osteoporosis risk did not overlap regions where many TFs were bound as shown in publicly available ChIP-seq profiles (49) (Fig. 3G and Supplementary Material, Table S8). However, cancer cell lines (and not ostb or adipocytes) were the main type of cells used for the ChIP-seq, and, in these types of studies, only a very small fraction of known TFs were assayed.

**Figure 3.**
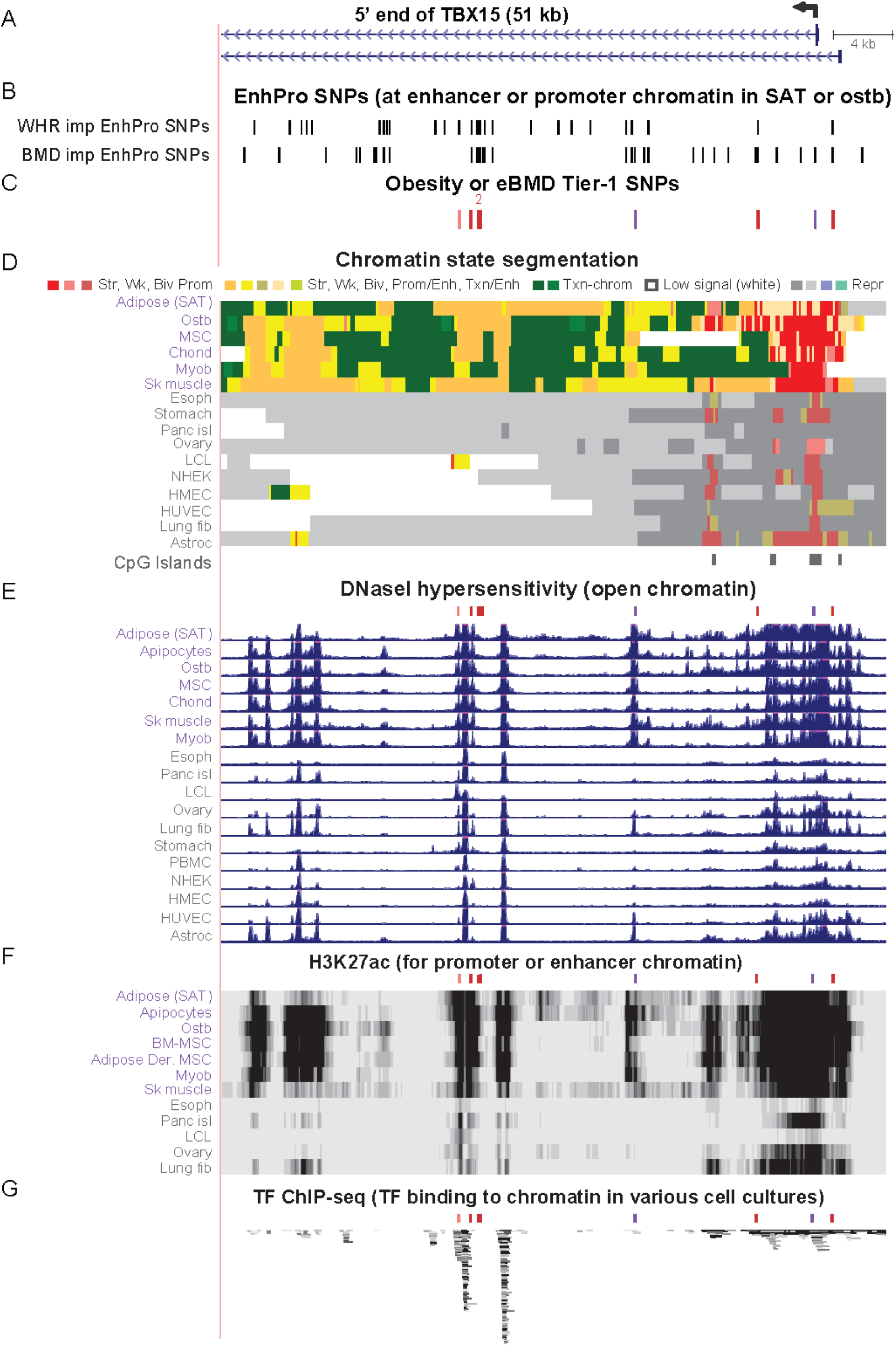
The 5’ end of *TBX15* contains all eight Tier-1 SNPs. (A) *TBX15*’s alternative first exons, part of intron 1 (chr1:119,484,631-119,535,630), and its main TSS (broken arrow). (B) WHR_adjBMI_ and eBMD imputed SNPs that are EnhPro SNPs. (C) The eight EnhPro SNPs that qualify as Tier-1 SNPs; “2” indicates two adjacent Tier-1 SNPs. (D) Chromatin state segmentation as in previous figures; Panc isl, pancreatic islets; Astroc, cultured astrocytes; CpG islands, CpG-rich regions as defined at the UCSC Genome Browser. (E) DNaseI hypersensitivity, vertical viewing range 0-10. (F) H3K27ac signal; vertical viewing range 0-4. (G) Binding of miscellaneous TFs to this region in a variety of cell cultures but not including ostb or adipocytes (see Supplementary Material, Table S8). Colored tick marks indicate positions of Tier-1 SNPs.

The trait-increasing allele (the allele associated with increasing WHR_adjBMI_, WC_adjBMI_, BMI, or eBMD) for six of the Tier-1 SNPs was the Alt allele and, importantly, the five Tier-1 SNPs, that were associated with obesity traits as well as eBMD, had the same trait-increasing allele for both traits (Table 2). With respect to gender differences, rs984222 and all obesity-associated index SNPs that are in high LD (r^2^ > 0.8) with it had much lower p-values for their association with obesity-related traits in women than in men (EUR), although the association for men was still highly significant (Supplementary Material, Table S1). We also noted similar gender directionality for Tier-1 SNPs rs1106529 and rs10923711 and the missense variant rs61730011. In contrast, rs9659323 and the other obesity- or eBMD-associated index SNPs that are in high LD with it exhibited lower p-values for their association with both traits in men than in women (EUR).

### Bayesian fine-mapping of *TBX15/WARS2* GWAS SNPs was not informative for causal SNP selection

We compared the results of using our criteria for identifying Tier-1 SNPs from WHR_adjBMI_ and eBMD GWAS to the results from fine-mapping the same imputed SNP data using the PAINTOR program (18) in baseline analyses. We found only two WHR_adjBMI_ SNPs with a posterior probability appreciably greater than 0, and these had a posterior probability of 1.0 and are in perfect LD with the above-described exon-3 missense variant, rs10494217 (Supplementary Material, Table S9). Only 11 eBMD SNPs had appreciable posterior probabilities greater than 0.10 and the highest was 0.52. These 11 eBMD SNPs were either in repressed chromatin in the intergenic region downstream of the 3’ end of *TBX15* or in intron 5 or 6 of this gene. Moreover, the results were almost identical when functional annotations for DHS overlap, chromatin state, and H3K27ac peak overlap at the SNPs in SAT or ostb were used in the fine-mapping.

### Three obesity- or osteoporosis-risk Tier-1 SNPs are in promoter or mixed promoter/enhancer chromatin near the 5’ end of *TBX15* in SAT and osteoblasts

Three of the Tier-1 obesity or eBMD SNPs (rs12742627, rs961470, and rs1106529) are within a 6-kb region spanning the main TSS for *TBX15* (Fig. 4B). The obesity and eBMD SNP rs1106529, which is 1 kb upstream of this TSS, was embedded in promoter chromatin in SAT or mixed promoter/enhancer-type chromatin in adipocytes and ostb. It has only two high-LD proxies, both of which are in intergenic repressed chromatin in adipose tissue and adipocytes (Supplementary Material, Table S4). The trait-increasing, Alt allele of rs1106529, but not the Ref allele, is predicted to overlap FOXO1, FOXC2, and SOX4 TFBSs (Table 2 and Supplementary Material, Table S7). These FOX family proteins are associated with both obesity and bone formation (Supplementary Material, Table S7) and the loss of SOX4 can cause osteopenia in mice (50) providing further support for rs1106529 being a strong candidate for a causal regulatory SNP related to both obesity traits and eBMD.

**Figure 4.**
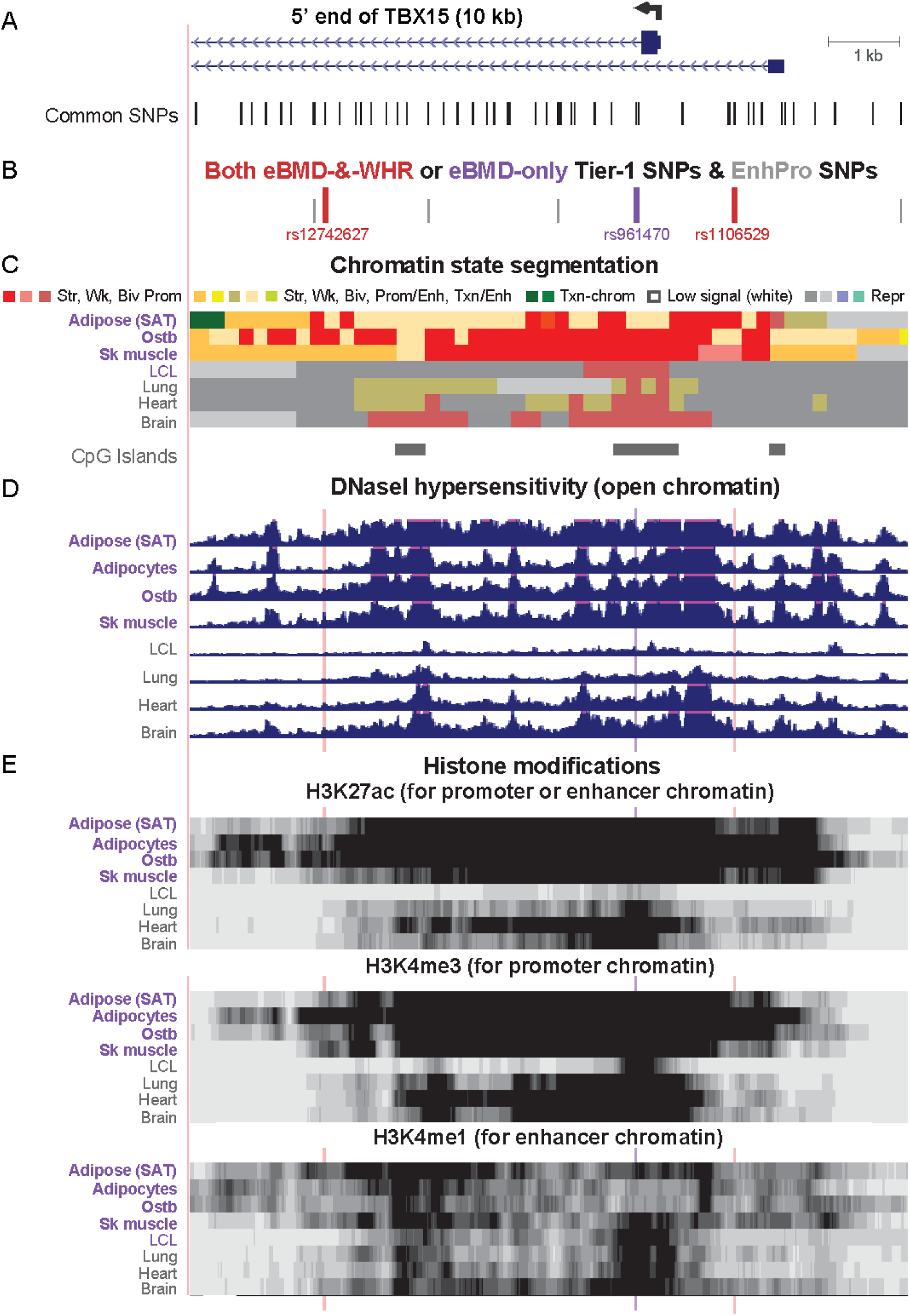
Zoomed in view of three Tier-1 SNPs near the main TSS of *TBX15*. (A) The common SNPs (version 151) at the 5’ end of *TBX15* (chr1:119,523,915-119,533,914). (B) The eBMD-and-WHR_adjBMI_ or eBMD-only Tier-1 SNPs are shown in red or purple, respectively; EnhPro SNPs that did not meet all the criteria for Tier-1 SNPs are denoted by short gray bars. (C), (D), and (E) as in Fig. 3 except that the vertical viewing range for DNaseI hypersensitivity was 0-7 and H3K4me3 and H3K4me1 are shown in addition to H3K27ac. Highlighting denotes the positions of Tier-1 SNPs.

The next two downstream Tier-1 SNPs, rs961470 (from eBMD GWAS) and rs12742627 (from eBMD and WHR_adjBMI_ GWAS) are in near-perfect LD with each other (Supplementary Material, Fig. S2A) and are located in intron 1. They are part of an extended TSS-downstream promoter or mixed promoter/enhancer chromatin region containing many DHS in SAT, adipocytes, ostb, MSC, and skeletal muscle but also in several tissue or cell types not expressing *TBX15* (Fig. 4A-D). However, the H3K27ac signals at and near these SNPs were much stronger or covered more of the chromatin in these expressing tissues than in tissues with little or no *TBX15* mRNA (Fig. 4E). The strongest epigenetic discriminator between the expressing and non-expressing tissues and cell cultures in the gene region was the chromatin state (Fig. 4C).

Only the trait-decreasing, Alt allele of rs961470 is predicted to bind to Nuclear Factor I-C (NFIC) and to SMAD3 (Table 2). Both of these TFs are implicated in regulating bone biology but while SMAD3 is generally associated with transcription activation, NFIC can be either a transcription repressor or activator (Supplementary Material, Table S7). For rs12742627, it is specifically the trait-increasing, Ref allele that is part of a predicted TFBS for E2F1, E2F4 (which are involved in normal bone formation) as well as E2F6, which is an inhibitor of E2F-dependent transcription, and TFDP1, which can heterodimerize with them (Supplementary Material, Table S7). Support for rs12742627 being embedded in an E2F family binding site comes from combined ChIP-seq data and TFBS predictions (UniBind, (48)), which demonstrated that E2F6 was bound to this sequence in human embryonic stem cells (ESC); neither ostb nor adipocytes were similarly tested.

### Three obesity- or osteoporosis-risk Tier-1 SNPs are in enhancer chromatin in a 0.7-kb cluster at *TBX15* intron 1

Three obesity-and-eBMD Tier-1 SNPs (rs984222, rs9659323, and rs1409157) are clustered within 0.7 kb with an additional WHR_adjBMI_ Tier-1 SNP (rs10923711) 1 kb further downstream in *TBX15* intron 1 (Fig. 5B). All four SNPs reside in strong enhancer chromatin in SAT, adipocytes, ostb, chond and MSC (Fig. 5C). Two of them, rs984222 and rs1409157 (which are in perfect LD), were among the seven eQTLs found for *TBX15* in SAT (Fig. 5B, lollipops) according to GTEx data (33) (Supplementary Material, Table S10). Although the p-values for these SAT eQTLs for *TBX15* are only in the range of about 1 to 3 x 10^-5^, all of them are also eQTLs for VAT (p = 1 x 10^-7^ to 7 x 10^-6^).

**Figure 5.**
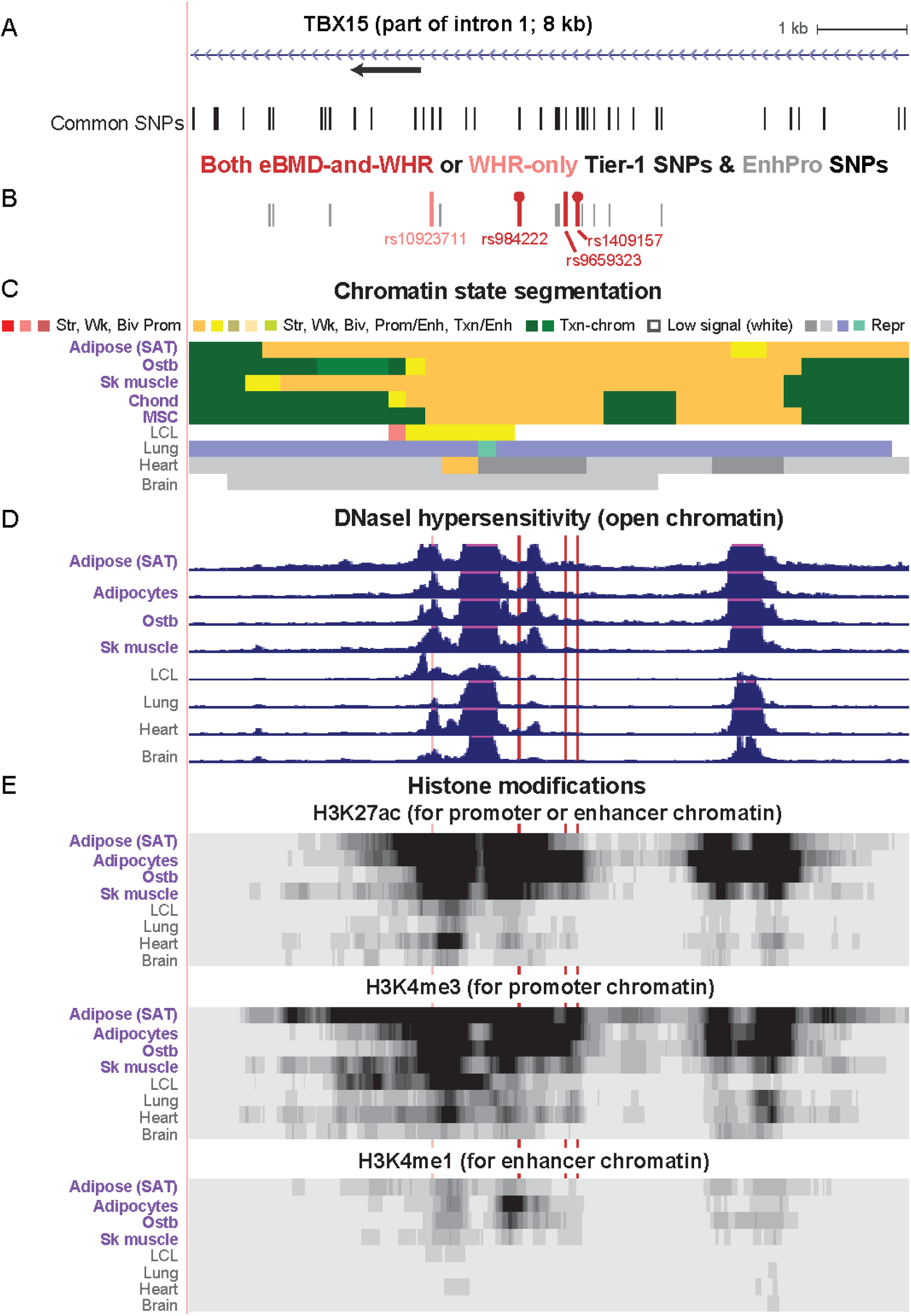
Zoomed in view of four Tier-1 SNPs in the middle of *TBX15* intron 1. (A) The common SNPs in this subregion of intron 1 of *TBX15* (chr1:119,500,183-119,508,182). (B-E) EnhPro SNPs and the subset of EnhPro SNPs that are designated Tier-1 SNPs, chromatin state segmentation, DNaseI hypersensitivity, and histone modifications as in Fig. 4 except that the CpG island track is not shown due to there being no CpG islands in this region. In panel B, lollipops denote the two *TBX15* eQTLs for SAT.

The trait-increasing, Alt alleles for rs984222 and rs1409157 are predicted to disrupt binding of MZF1, ZNF281, GATA2, or GATA6 TFs (Table 2). These TFs are implicated in regulating the differentiation in the adipose and/or bone lineages (Supplementary Table S7). Rs9659323 is at a predicted allele-specific TFBS for numerous members of the AP-1 (Fos/Jun) family as well as for MAF/NFE2 heterodimers. AP1 family members are involved in ostb maturation, formation of bone matrix proteins in ostb and chond, and adipocyte differentiation and, like the above-described TFs, all can act as transcriptional activators (Supplementary Material, Table S7). Rs9659323 borders a 142-bp DNA repeat (short interspersed nuclear element, SINE) while rs984222 is in the middle of a 185-bp DNA repeat (long interspersed nuclear element, LINE; Supplementary Material, Table S6). This overlap of repeats might be functionally important because some transcription elements can arise by exaptation of SINEs and LINEs (51). Although, rs9659323 and rs984222/rs1409157 are only in moderate LD (r^2^ = 0.45, EUR; Fig. 6A), the frequency of the haplotype with trait-increasing alleles for all three of these SNPs is considerable, ∼41% of the EUR population (Fig. 6B).

**Figure 6.**
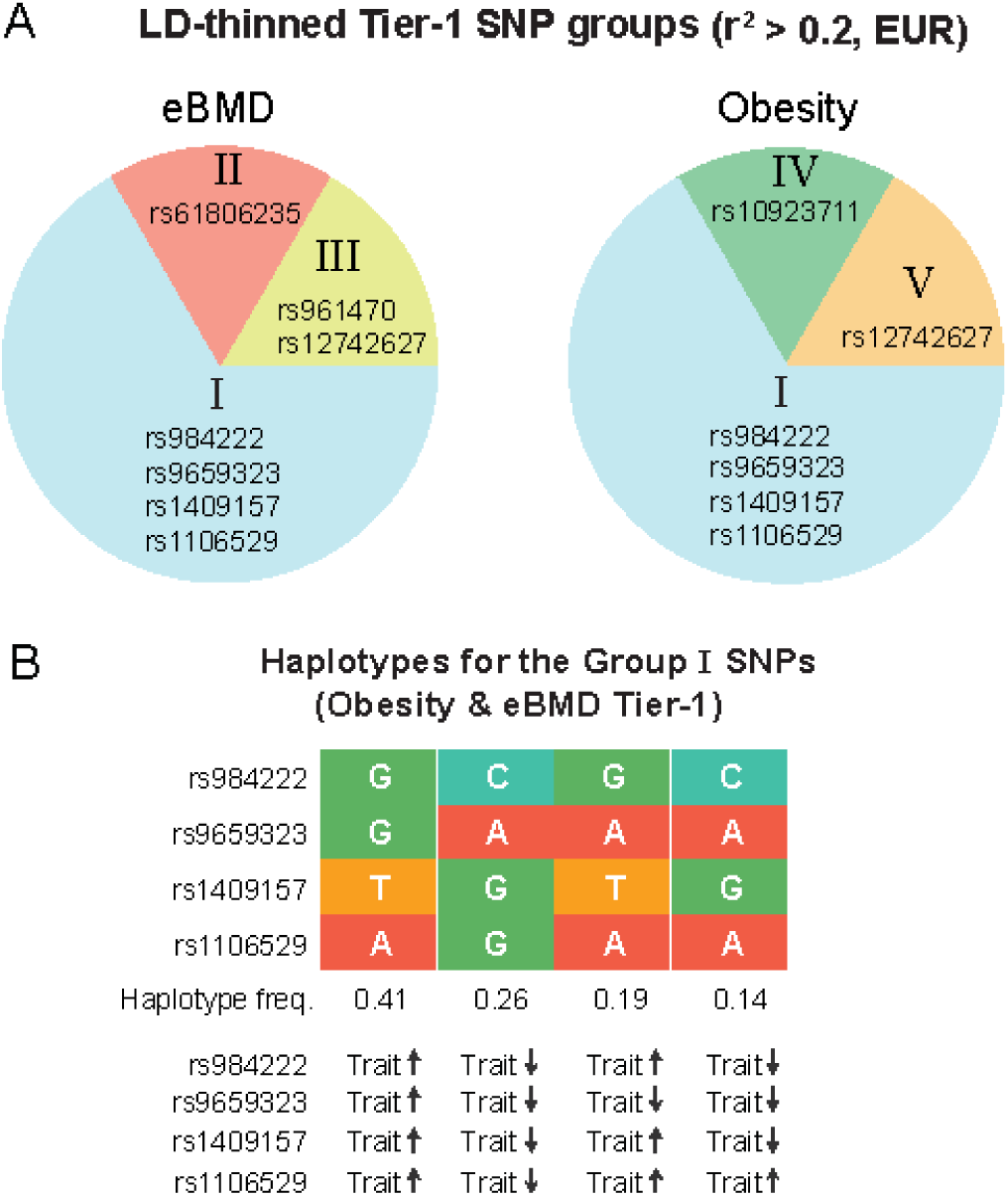
The genetic architecture of the Tier-1 SNPs at *TBX15* suggests that there are multiple causal regulatory SNPs for obesity risk and for osteoporosis risk. (A) The SNPclip program at LDlink (52) was used to trim the Tier-1 SNPs into groups with r^2^<0.2 (EUR); the SNP members of the resulting groups are shown in pie charts. (B) The haplotypes for trimmed Group I are shown with designations of trait-increasing (trait ↑) or trait-decreasing (trait ↓) alleles; the trait-increasing alleles from obesity GWAS are the risk alleles but for eBMD GWAS they are the non-risk alleles because of the osteoporosis-protective effects of high BMD.

The most distal Tier-1 SNP in this region, the obesity-related rs10923711, was in a moderately strong DHS in SAT and adipocytes (Figs. 3E and 5D) and is predicted to bind with allelic specificity (for the Ref allele) to ETS2, ETV5, and ELK1, two of which have been described as being related to adipose biology (Supplementary Material, Table S7). Although rs10923711 is the closest Tier-1 SNP to the *TBX15* exon-3 missense SNP rs10923711, its LD with that SNP or the exon 8 missense SNP (rs61730011) was very weak (r^2^ = 0.02 or 0.05, EUR). It was in low LD (r^2^ = 0.16 or 0.17) with the three clustered Tier-1 SNPs (rs984222, rs9659323, and rs1409157) that are ∼1 kb distant (Supplementary Material, Fig. S2), and so it was not in the same LD-thinned variant group (52) as they (Table 2 and Fig. 6A, right). This is in contrast to rs1106529, which is 27-kb downstream in the promoter region but in the same LD-thinned group as the 0.7-kb cluster of three Tier-1 SNPs.

### Obesity or eBMD SNPs overlapping *WARS2* are probably only proxies for *TBX15* intron-1 SNPs

Two obesity-related index SNPs, rs17023223 and rs2645294 (Table S1), are found in the gene body of the broadly expressed neighbor to *TBX15* namely, *WARS2* (Fig. 2C, black triangles). Located in the 3’ UTR of *WARS2,* rs2645294 is one of the seven *TBX15*-specific eQTLs in SAT (Supplementary Material, Table S10) while rs17023223 is embedded in weak enhancer chromatin at intron 1 of *WARS2* in SAT. Neither of these SNPs nor their high LD (r^2^ > 0.8) proxies met the definition of a Tier-1 SNP. However, they are in moderately high LD with the three Tier-1 SNPs in the 0.7-kb cluster in intron 1 of *TBX15*. The LD values for rs2645294 with rs984222 or rs1409157 and for rs17023223 with rs9659323 are r^2^ = 0.76 and 0.71 (EUR), respectively. Moreover, just as there was a much stronger association of WHR_adjBMI_ or WC_adjBMI_ with women than men for index SNP rs984222, so index SNP rs2645294 showed a strong bias for women in WHR_adjBMI_ while that was not the case for rs17023223 or rs9659323 (Supplementary Material, Table S1). Therefore, the *WARS*2 index SNPs are probably associated with obesity risk through their linkage to several *TBX15* Tier-1 SNPs. Nonetheless, the presence of many *TBX15* eQTLs for VAT in the *WARS2-*adjacent non-coding RNA (ncRNA) gene *LOC101929147* and many *WARS2*-specific eQTLs for SAT (and VAT) embedded in *TBX15* or the downstream gene desert (Fig. 2D) suggests that this gene neighborhood is part of large unit of higher-order chromatin structure with biologically important interactions. This interpretation is consistent with the finding of only few and weak sites for CCCTC-binding factor (CTCF, a chromatin conformation-altering protein) in this gene neighborhood in various cell lines, including ostb (Fig. 2G).

### *TBX18*, a closely related paralog of *TBX15*, is associated with two Tier-1 SNPs for eBMD or obesity traits

*TBX18* is structurally and functionally related to *TBX15* (26) but located on a different chromosome (Supplementary Material, Fig. S3). Like *TBX15, TBX18* is preferentially expressed in ostb, MSC, chond, SAT, and adipocytes and shows much lower expression in VAT (Supplementary Material, Table S2). Using the criteria described for evaluating the most likely causal SNPs in *TBX15*, we found an eBMD Tier-1 SNP, rs61543455 (p = 1.2 x 10^-10^ (27)) for *TBX18* in the gene’s far-downstream region (Supplementary Material, Figs. S3 and S4). Rs61543455 overlaps strong enhancer chromatin and a strong DHS in ostb and MSC. It is located at the 5’ end of an ncRNA gene (*ENST00000454172*), the nearest gene upstream of *TBX18*, which has a similar transcription profile in tissues to that of *TBX18* (Supplementary Material, Fig. S3G). This SNP overlaps allele-specific TFBS for FOXO1, SOX4 and FOXP1 TFs. The first two of these TFs were also predicted to bind in an allele-specific manner to the *TBX15*-upstream Tier-1 SNP (rs1106529; Table 2) and can positively regulate osteogenesis (Supplementary Material, Table S7). In addition, the nearby eBMD SNP rs17792917 overlaps strong enhancer chromatin and a DHS in chond and weak promoter chromatin but not a DHS in ostb. It is located 1 kb upstream of the TSS of *ENST00000454172* and is predicted to be an allele-specific TFBS for ARID5A, a chondrocyte-associated TF (53). Therefore, these two SNPs might impact osteoporosis risk by influencing *TBX18* expression in ostb or chond by an indirect effect on transcription of an upstream ncRNA gene. No WHR_adjBMI_ Tier-1 SNPs were seen near *TBX18*.

### Discussion

Using in-depth manual curation, we prioritized eight variants as best candidates for regulatory causal SNPs (Tier-1 SNPs; Table 2) from 692 *TBX15/WARS2*-region SNPs that are associated with obesity traits or eBMD. Only two of these Tier-1 SNPs, rs984222 and rs1106529, had been described previously as possible causal SNPs for obesity (9, 10, 20, 54, 55), and none for osteoporosis. For selection of the Tier-1 SNPs, we made the following assumptions. First, a regulatory SNP is likely to be in or near a gene that is preferentially expressed in the phenotype-relevant tissue. Second, it should overlap enhancer or promoter chromatin selectively in this trait-related tissue (10, 14, 15, 19, 54). Third, it should overlap a DHS in this tissue because a DHS is indicative of TF binding (56). Last, the SNP should be within a stringently predicted allele-specific TFBS whose corresponding TF is expressed in the phenotype-relevant tissue (57). One caveat to these assumptions is that, besides the SAT and ostb trait-associated cell populations which we studied, there may be other cell types through which regulatory SNPs in the *TBX15/WARS2* neighborhood affect obesity or osteoporosis risk, e.g., bone-degrading osteoclasts (58), for which detailed *TBX15* expression and epigenetic data are not publicly available. Another caveat is that an obesity- and/or osteoporosis-risk SNP that does not overlap a single predicted allele-specific TFBS might assert its effect on transcription through interactions with multiple TF complexes (59). Lastly, there may be local SNP-related transcription-regulatory potential that is not captured by the studied H3 modifications or CTCF binding (Figs. 2G and 5E).

All eight Tier-1 SNPs were within or immediately upstream of *TBX15* even though the GWAS-identified SNPs extended over a 433-kb region. In this region, a *TBX15*-upstream ncRNA gene, *LOC105378933,* has an expression profile similar to that of *TBX15* (Table 1). However, none of the EnhPro SNPs within or upstream of this ncRNA gene met the above-described criteria for Tier-1 SNPs. The nearest protein-encoding gene to *TBX15, WARS2,* was reported to have a significant association of its SAT and VAT RNA levels (by qRT-PCR) with obesity (20). In addition, the distribution of GTEx-determined eQTLs for SAT and VAT (Fig. 2) suggests that the upstream and downstream neighborhood of *TBX15/WARS2* is a unit of higher-order chromatin structure. *WARS2* encodes a broadly expressed, mitochondrial tryptophanyl tRNA synthetase, which could impact metabolism in adipose tissue through effects on mitochondria. Nonetheless, it is likely that the two index *WARS2*-overlapping obesity-trait associated SNPs and their enhancer chromatin-overlapping proxies are probably associated with obesity traits in GWAS just because they are in high LD with Tier-1 SNPs in the 0.7-kb cluster at *TBX15* intron 1.

All the obesity Tier-1 SNPs and three of the eBMD Tier-1 SNPs associated with *TBX1*5 were embedded in the gene’s SAT or ostb super-enhancer. Such large enhancers play a major role in tissue-specific transcription regulation of developmental genes (39, 60). Most of the TFs predicted to bind in an allele-specific manner to Tier-1 SNP-containing sequences have known biological associations with adipose or bone biology (Tables 2 and Supplementary Material, Table S7). In contrast to the eight candidate causal SNPs found from our manual curation, no good candidate regulatory SNPs for modulating obesity risk and only poor candidates for eBMD risk were found when we evaluated large sets of imputed SNPs in the *TBX15/WARS2* region by fine-mapping using the PAINTOR program (61). Manual curation of epigenetic and transcriptomic data revealed that the Tier-1 SNPs often overlapped only small DHS in the tissue or cell type of interest. Small DHS can denote TF binding *in cis*, just as for large DHS (62), although the smaller peaks probably indicate fewer TFs bound to that subregion. Our stringent manual curation of allelic TFBS overlapping SNPs eliminated as poor matches some SNP-containing sequences that TFBS matching programs, including TRANSFAC and JASPAR, rated as good matches. In addition, manual inspection of transcriptomics and chromatin state segmentation databases produced evidence for a small transcript (*ENST00000449873.1*) initiated within the 3’ portion of the canonical *TBX15* gene (Fig. 1) that was almost the only transcript in liver. Surprisingly, according to the GTEx database (33), this truncated transcript comprised about half of the *TBX15*-derived transcripts in SAT. Although this transcript was assigned protein-coding potential by ENSEMBL, its biological role is unknown, and it does not include sequences specifying the *TBX15* DNA-binding domain. Given the overlap of four of the Tier-1 SNPs with enhancer chromatin in liver, such a transcript might contribute to the examined risk associations of SNPs in this region.

TBX15 can heterodimerize with the closely related TBX18 to form potent repressors working in conjunction with Groucho (TLE) corepressors (63). Remarkably, our analysis of eBMD GWAS data (64) indicates that the *TBX18* gene is also associated with eBMD. This is consistent with the partial overlap of functions and high amino acid sequence homology between TBX15 and TBX18 proteins (26, 65).

Our analysis of GWAS data highlighting *TBX15* as both an osteoporosis- and obesity-related gene harboring risk SNPs is consistent with the importance to bone development of TBX15 (25) and the relationships of this TF to adipose biology (20, 22, 23, 34, 66, 67). It is also in accord with the interrelationships in the development of bone and adipose tissue (68) and the preferential expression of *TBX15* in both adult-derived ostb and SAT. The high and tissue-specific expression of *TBX15* in SAT suggests that some of its functions in the adipose lineage involve homeostasis or responses to physiological changes. *TBX15* is expressed in a fat depot-specific manner (20, 23, 66) despite heterogeneous expression among the SAT-derived adipocytes and among preadipocytes expressing adipose-lineage markers (66). It is also preferentially expressed in MSC (which can give rise to ostb) and chond (which are in an alternative pathway to bone formation) (23, 66, 69). There is also evidence for the involvement of TBX15 in modulating white and brown/brite adipocyte differentiation; mitochondrial function and oxidative or glycolytic metabolism in adipose cells; and cold-adapted adipose metabolism among the Inuit (22, 23, 34, 66, 67). In addition, this TF has a well-established role in the embryonic development of many parts of the skeleton (25, 26).

Whether *TBX15* has a positive or negative effect on adiposity is controversial. Gesta et al. (24) reported that five-fold overexpression of *TBX15* impaired differentiation in a preadipocyte cell line and reduced mitochondrial mass and triglyceride accumulation. However, in a previous study the same group reported a positive relationship between WHR or BMI increases and increases in qRT-PCR-determined *TBX15* mRNA levels in human abdominal SAT but the opposite relationship in VAT (70). Sun et al. found that homozygous knockout of *Tbx15* targeted to adipose tissue in mice was associated with increased fat mass when the mice were fed high-fat diets (71) suggesting that TBX15 levels are negatively correlated with obesity. They observed impaired adipocyte browning in inguinal adipose (SAT) upon cold challenge, which they related to the weight gain. In contrast, Singh et al. reported that *TBX15* null mice were leaner than their control littermates in addition to having reduced bone size and muscle mass (25). Gburcik et al. (23) found that knock-down of *Tbx15* expression decreased transcription of white and brown/brite adipogenic marker genes in short-term differentiating cultures of preadipocytes from inguinal or brown adipose depots. From adipose-specific *Tbx15* heterozygous mouse knockouts, Ejarque et al. (22) observed no change in differentiation in *TBX15* knockdown experiments on preadipocytes but reported higher levels of TBX15 protein (by immunoblotting) in SAT from obese vs. lean individuals.

A positive correlation of *TBX15* expression with obesity is suggested by the finding that rs984222 (one of our Tier-1 SNPs) is an eQTL whose risk allele is significantly and positively associated with *TBX15* expression in SAT (33) and in the much lower-expressing VAT (21, 33). In considering the relationship of *TBX15* to obesity, it should be noted that obesity is associated with increases in both SAT and VAT depots (5). The relationship of *TBX15* to obesity is probably complicated by the depot-dependent levels of *TBX15* expression. Importantly, GWAS-derived SNPs in *TBX15* were derived from associations with different obesity traits, WC_adjBMI_ and BMI as well as in WHR_adjBMI_ (Supplementary Table S1). A further complication in understanding the role of *TBX15* in obesity from experimental studies is that fine-tuning changes in *TBX15* activity, like the very small alterations in expression expected from GWAS regulatory SNPs, might have different directional effects on *TBX15* functionality than large experimentally induced changes.

The direction of the relationship between *TBX15* expression and bone health is much clearer. Homozygous inactivating mutations in *TBX15/Tbx15* in humans (Cousin syndrome) and mice (double knockout and Droopy ear homozygous mutant) are linked predominantly to multiple skeletal abnormalities (26, 72). Moreover, studies of embryogenesis in normal and *Tbx15* null mutant mice implicate Tbx15 protein in controlling development of bone throughout the body during embryogenesis possibly by up-regulating the number of early bone precursor cells (MSC and chondrocytes) (25). Interrelationships between the bone and adipose lineages include the findings that MSC and mesenchymal progenitor cells can to give rise to either ostb or adipocytes depending on the cellular milieu (2, 28, 36). Postnatally, similar steady-state levels of *TBX15* mRNA and shared promoter and enhancer chromatin were seen in ostb, chond and MSC (Fig. 1), which suggests that *TBX15* Tier-1 SNPs might play a role in modulating BMD risk in all three of these cell types.

Part of the biological effect of *TBX15*-associated Tier-1 SNPs on obesity traits and eBMD might occur through an anti-apoptotic mechanism in adipose and bone precursor cells (ostb, MSC, and chond) and in the corresponding mature cells (adipocytes and osteocytes) prenatally and postnatally. A common function for TBX15 in adipose and bone biology might explain our remarkable finding that each of the five Tier-1 SNPs that came from both obesity trait and eBMD GWAS had the same trait-increasing allele for obesity as for eBMD. TBX15 was reported to have an anti-apoptotic role in examined thyroid cancer cell lines (17). Apoptosis can alter adipose tissue both at the adipocyte and preadipocyte levels in normal stressed or unstressed tissue (3, 73). It is also a major shaper of prenatal and postnatal bone biology by affecting numbers of bone progenitor cells (MSC/chond/ostb), bone-degradative osteoclasts, and osteocytes (36) although there has been only very little examination (25) of the effects of *TBX15* on apoptosis in bone or adipose cells or progenitors. Other ways that TBX15 might impact obesity risk and eBMD is by increasing progenitor cell numbers through a mechanism involving cell cycling (25, 74) or cell differentiation (24). In the case of either decreased cell death or increased cell cycling, the result could be strengthening bone (increased eBMD correlated with good health) or increased adiposity (correlated with poorer health when in VAT or deep superficial abdominal SAT (5)).

We propose that multiple SNPs, rather than a single one, in the gene body or promoter region of *TBX15* fine-tune regulation of this gene’s expression in target tissues and thereby alter the genetic risk of obesity and osteoporosis. This conclusion is in accord with our finding that the obesity or eBMD Tier-1 SNPs each fell into three separate thinned variant groups based on genetic architecture (Fig. 6), each of which had their SNPs in enhancer or promoter chromatin found preferentially in SAT or ostb/MSC/chond. The two missense SNPs in exons 3 and 8 constitute two additional thinned variant groups. Most of the TFs corresponding to the predicted allele-specific TFBS can either repress or activate transcription depending on the chromatin context and were found to have special roles in adipose or bone biology (Supplementary Material, Table S7). One of the thinned groups of SNPs (Group I; Fig. 6B) contains Tier-1 SNPs from both obesity-trait and eBMD GWAS. The four Tier-1 SNPs in this group (rs984222, rs9659323, rs1409157 and rs1106529), three of which are clustered within 0.7 kb in a SAT super-enhancer and in ostb enhancer chromatin in *TBX15,* might act together in a Alt-allele risk haplotype that has a frequency of about 0.41 (EUR; Fig. 6B) (15). Nonetheless, without experimental assays, we cannot discount the possibility that these four SNPs are correlated just because of LD irrespective of biological cooperation. In conclusion, our in-depth, manually-curated bioinformatics analysis of hundreds of GWAS-derived SNPs from the *TBX15/WARS2* region led to only eight prioritized regulatory SNPs as prime risk candidates for obesity and/or osteoporosis risk., all of which were at the 5’ end of *TBX15* and apparently acting independently of two *TBX15* missense SNPs.

## Methods

### GWAS-derived SNPs and LD analysis

*TBX15/WARS2* and *TBX18* SNPs that were significantly (p < 5 x 10^-8^) associated with WHR_adjBMI_, WC_adjBMI_, BMI and eBMD were retrieved from the NHGRI-EBI GWAS Catalog (downloaded August, 2019; (37)) and termed as index SNPs. The index SNPs were expanded to a set of proxy SNPs (r^2^ ≥ 0.8 in a 1-Mb window, EUR population from the 1000 Genome Project Phase 3; (75)) by using PLINK v1.9 (76). In addition, we retrieved summary statistics datasets for WHR_adjBMI_ (38) and eBMD (27), which are the largest publicly available summary statistics for these traits, and extracted imputed significant SNPs (p < 6.6 x 10^-9^) for *TBX15* (chr1:118,717,862-119,916,299) and for *TBX18* (chr6:84,921,843-86,171,855). For subdividing the Tier-1 SNPs into thinned LD groups (pairwise r^2^ > 0.2), EUR, we used the SNPclip function of the LDlink suite (52). From LDlink, LDmatrix (EUR) was used for the heat map of LD and LDhap for the haplotype analysis.

### Transcriptomics

Tissue RNA-seq data were from Genotype-Tissue Expression (GTEx) RNA-seq median gene expression levels (33) for 36 tissue types (Table S2; median TPM from hundreds of samples for each tissue type). The GTEx SAT samples were from beneath the leg’s skin and VAT samples from the parietal peritoneum. For cell cultures, we used total RNA-seq data (ENCODE/Cold Spring Harbor Lab) for 12 cell culture types (Table S3; average RPKM of technical duplicates) (77) rather than poly(A)^+^ RNA because this was the only type of RNA-seq available for ostb, chond, and MSC; in Fig. 1, the vertical viewing range was 0 – 200. For RNA-seq, the ostb were from femoral bone of a 62 y male and a 56 y female; bone marrow-derived MSC (referred to as MSC) were from femoral bone marrow of a 57 y male and a 60 y female; chond from knee or femoral cartilage of a 64 y male and a 56 y female; preadipocytes were undifferentiated cells from white SAT pooled from two individuals. The exact origins of ostb and bone marrow-derived MSC for ENCODE and Roadmap samples is not known but chond were obtained by *in vitro* differentiation of bone marrow-derived MSC.

### Epigenetics

All epigenetic data are from Roadmap Project (14) and available at the UCSC Genome Browser (http://genome.ucsc.edu/) (35). Chromatin state assignments at the SNPs of interest came from the 18-state chromatin state segmentation track with promoter chromatin as states 1, 2 and 4, enhancer chromatin as states 7-11, and mixed promoter/enhancer chromatin as state 3. In figures, the color-coding of Roadmap for the promoter and enhancer chromatin states was slightly modified for clarity as shown in the figures. Obesity EnhPro SNPs were defined as obesity-trait GWAS-derived SNPs embedded in promoter or enhancer chromatin present in SAT and no more than two of the 11 other tissue types from among the following tissues: aorta, liver, pancreas, pancreas islet, lung, spleen, left ventricle of the heart, brain dorsolateral prefrontal cortex, CD14^+^ monocytes, peripheral blood mononuclear cells and psoas skeletal muscle. eBMD EnhPro SNPs are eBMD

GWAS-derived SNPs within promoter or enhancer chromatin in ostb but in no more than two of the following 12 types of cell cultures: lung fibroblasts, fetal lung fibroblasts (IMR90), keratinocytes and melanocytes from foreskin, mammary epithelial cells, myoblasts, umbilical vein endothelial cells, astrocytes, adult dermal fibroblasts and epidermal keratinocyte primary cell cultures or ESC (H1) and lymphoblastoid (GM12878) cell lines. The adipose tissue sample for Roadmap chromatin state segmentation and DNaseI hypersensitivity profiling is a mixture of five samples from females (49Y, 59Y, 41Y, 25Y and 81Y) (14). It was not designated as SAT in the Roadmap project but is in the ENCODE project (https://www.encodeproject.org/search/?type=Biosample). This assignment is confirmed by the extensive strong enhancer and promoter chromatin overlapping *TBX15* in this adipose sample (Fig. 1) and the high expression levels of *TBX15* in SAT samples and much lower levels in VAT samples (>300 samples each, Supplementary Material, Table S2). It is also consistent with the lack of promoter or enhancer chromatin for *WT1* (14, 35), a marker for VAT (4). For other comparisons, we used Roadmap epigenetic data from mixed bone marrow-derived MSC cultures from four individuals, adipocytes from bone marrow MSC (female), as well as the other tissues and cell types given in figures and tables. DHS for EnhPro SNPs were scored as narrow peaks (narrowPeaks from DNase-seq) for ostb, SAT, and adipocytes except for rs9659323 and rs1409157, for which the designation was based on visual assessment of DHS tracks using the UCSC Genome Browser.

### Allelic TFBS predictions

To determine whether EnhPro SNPs were embedded in predicted TFBS, we used primarily the UCSC Genome Browser track for JASPAR 2018 (35, 47) and TRANSFAC (44) programs and secondarily SNP2TFBS (46) and HaploReg v4.1 (45). Unlike the other three programs, JASPAR gives predictions only for the Ref allele. For JASPAR matches, we considered only TFs with scores ≥ 400 unless a perfect match was found in the literature for the examined binding site (see rs984222 and MFZ1; Supplementary Material, Table S7). For TRANSFAC predictions, we used a 15-b input sequence with the SNP in the middle and required matrix scores > 0.8 and core scores > 0.9 using the 2019.2/Vertebrate TFBS database and the settings for Only High Quality Matrices and Minimize False Positives. For HaploReg, we used only the “known” position weight matrix (PWM) data that gave a difference of ≥ 5 for the HaploReg Ref vs. Alt scores. For all the predicted TFBS, the corresponding sequence-specific DNA binding TF had to have a TPM of ≥ 2 in SAT or a moderate or strong signal in ostb in the RNA-seq track at the UCSC Genome Browser (see above). We used manual curation to retain only those predictions for which all of the conserved positions in the PWM had exact matches and no more than one base in a partly conserved position had only a partial match (≤ 5 fold difference in binding of the genomic base compared with the PWM-favored base). For determining that a TFBS overlapping at a SNP is likely to be allele-specific in its TF binding, we required that the disfavored allele had a position probability that was ≥5-fold lower than that of the favored allele in the PWM matrix. We checked all the Tier-1 SNPs for overlaps to experimentally determined TFBS in the UniBind and Factorbook tracks for TF-ChIP-seq in the UCSC Genome Browser (48, 49).

### Summary statistics imputation and fine-mapping analyses

We first augmented the GWAS summary statistics imputed SNPs from the sex combined WHR_adjBMI_ dataset of Pulit et al. (38) using ARDISS (Automatic Relevance Determination for Imputation of GWAS Summary Statistics) (78) and all their summary statistics-derived SNPs for chr1:119,369,719-119,803,112 (hg19) to obtain the maximum overlap of our obesity-related index/proxy SNPs with obesity imputed SNPs (77%). This enhanced imputation was only used for fine-mapping of the WHR_adjBMI_ SNPs. For the eBMD fine-mapping the imputed SNPs from Morris et al. (27), were used without further imputation because there was >70% overlap of eBMD index/proxy SNPs with these imputed SNPs. We, then, applied a Bayesian approach, PAINTOR (Probabilistic Annotation INtegraTOR, version 3.1)(61), which is an Expectation Maximization algorithm that integrates Z-scores for association between each SNP and the trait, LD structures, and functional annotations, to calculate the posterior probability for each SNP and prioritize the most plausible causal variants at a given locus. The 1000 Genome Project Phase 3 (EUR) (75) served as the reference genotypes for imputation and fine-mapping analyses. For functional annotations, we queried overlap of the SNPs with the following attributes: exonic DNA, 165 PAINTOR-provided TFBS annotations, Roadmap-derived enhancer and promoter chromatin segmentation state, and H3K27ac in ostb and SAT, DHS narrow peak in ostb and adipocytes, and CTCF binding sites in ostb.

## Supplementary Material

Four Figures are in a Word file and 10 Tables in 10 Excel files.

## Acknowledgements

We are very grateful to Dr. Bogdan Pasaniuc and Ms. Ruth Dolly Johnson for their generous help with the PAINTOR analyses.

