## Supplemental Figures for "Osteoporosis- and obesity-risk interrelationships: An epigenetic analysis of GWAS-derived SNPs at the developmental gene *TBX15*"

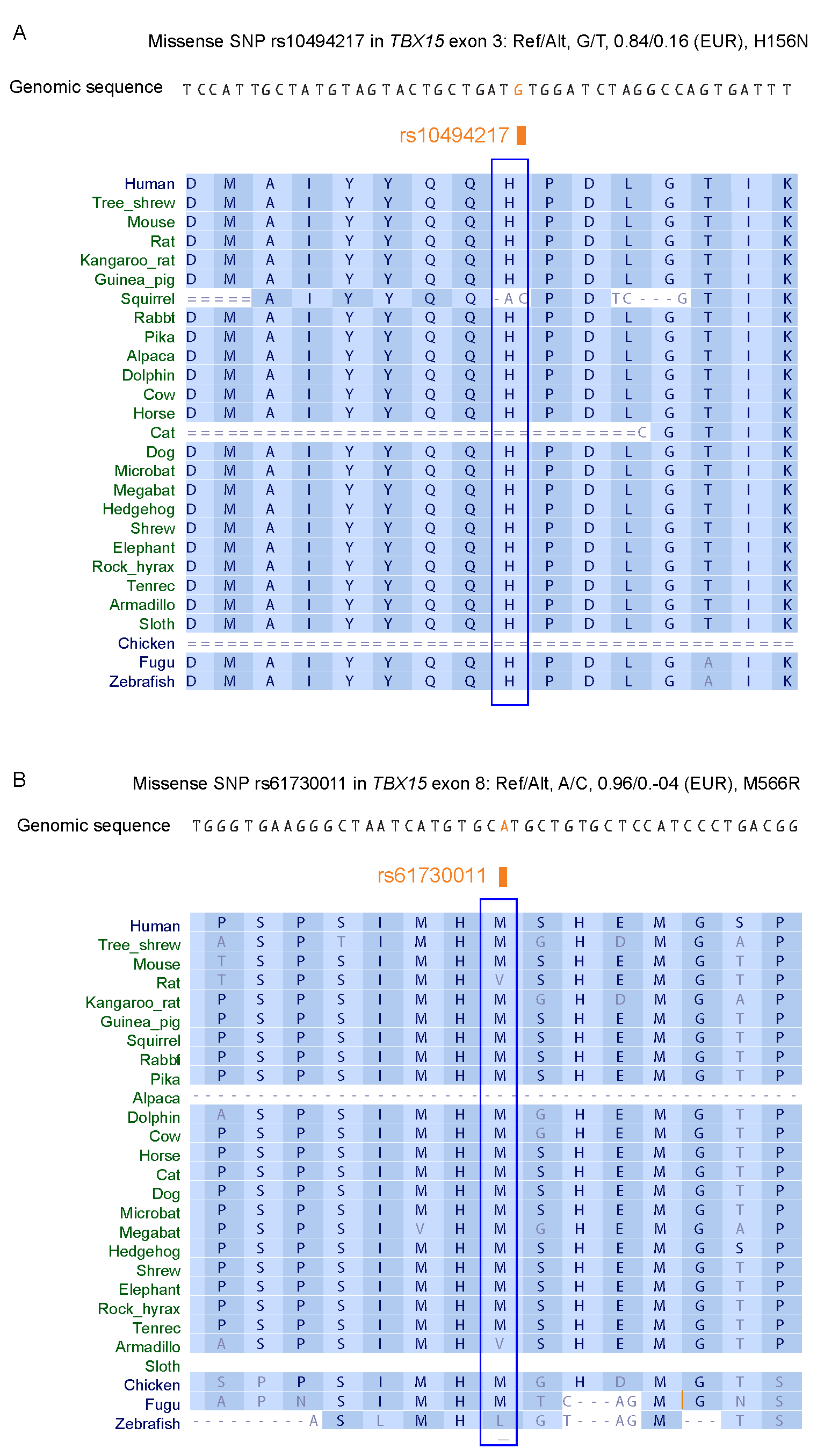
**Figure S1.** **Strong conservation of amino acid sequences near the two *TBX15* missense SNP alleles associated with obesity traits**. **(A)** Missense SNPs rs10494217 and **(B)** rs61730011, which are associated with WHR_adjBMI_ and eBMD or just with WHR_adjBMI_ , respectively (1, 2), are shown with the genomic sequence at these subregions in the human RefSeq genome (Ref allele). The corresponding codons in humans and several other vertebrates are aligned. The data are for chr1:119,469,163-119,469,208 and chr1:119,427,444-119,427,489 (hg19), respectively, and are from the Multiz alignment/Conservation 46-way track at the UCSC Genome Browser (3). Light gray font, amino acids that differ from those in humans.

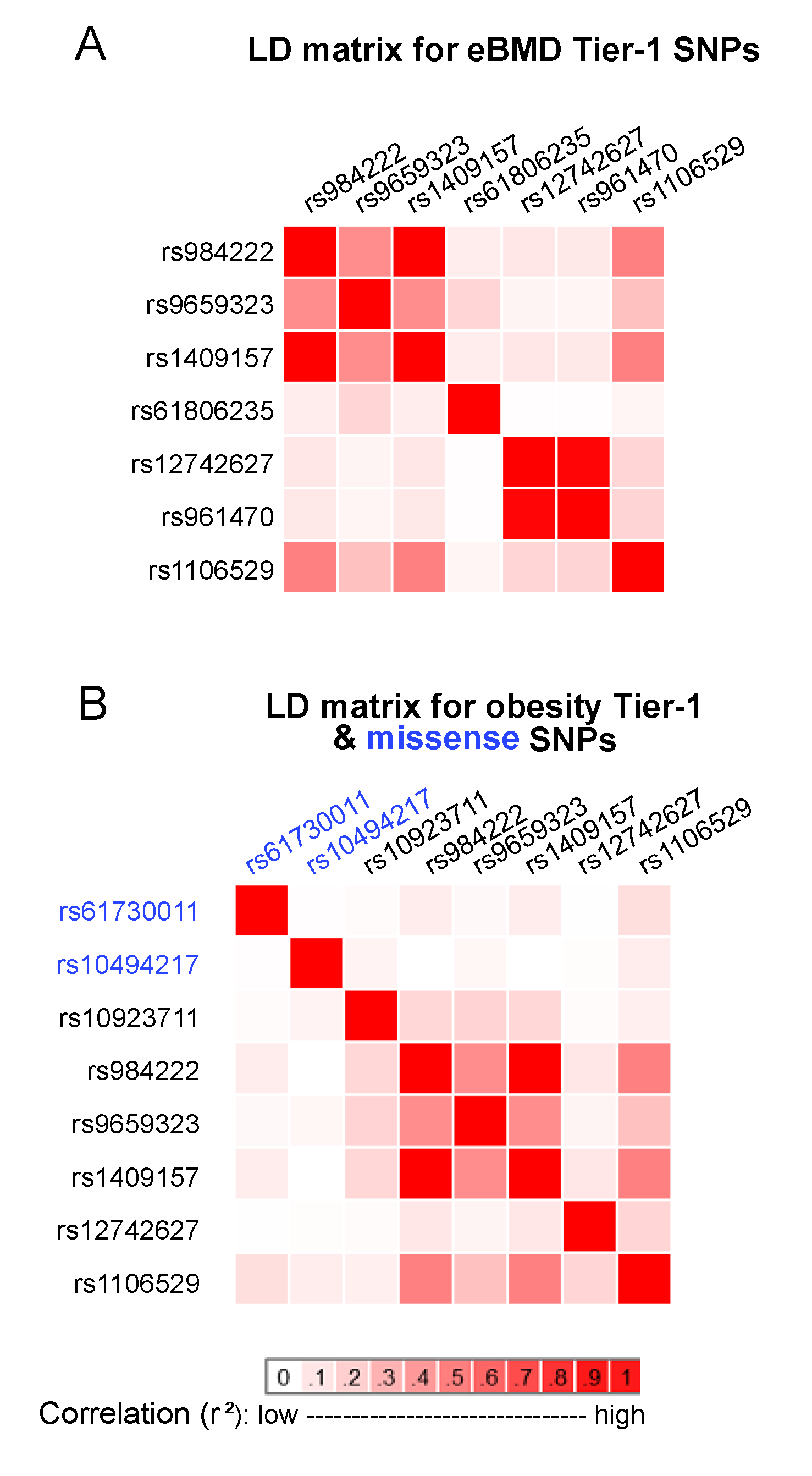
 **Figure S2. Linkage disequilibrium (LD) matrices for Tier-1 SNPs and missense SNPs linked to *TBX15* show that the Tier-1 and missense SNPs are not in appreciable LD but subgroups of the Tier-1 SNPs are**. LD matrices for **(A)** eBMD-linked Tier-1 SNPs and for **(B)** obesity-linked Tier-1 SNPs and the two missense SNPs (blue) for the European ancestry (EUR) population, as determined by the LDLink program (4). Rs984222 and rs1409157 are perfectly correlated as to LD (r^2^ = 1.0); rs12742627 and rs961470 are in near perfect LD (r^2^ = 0.98).

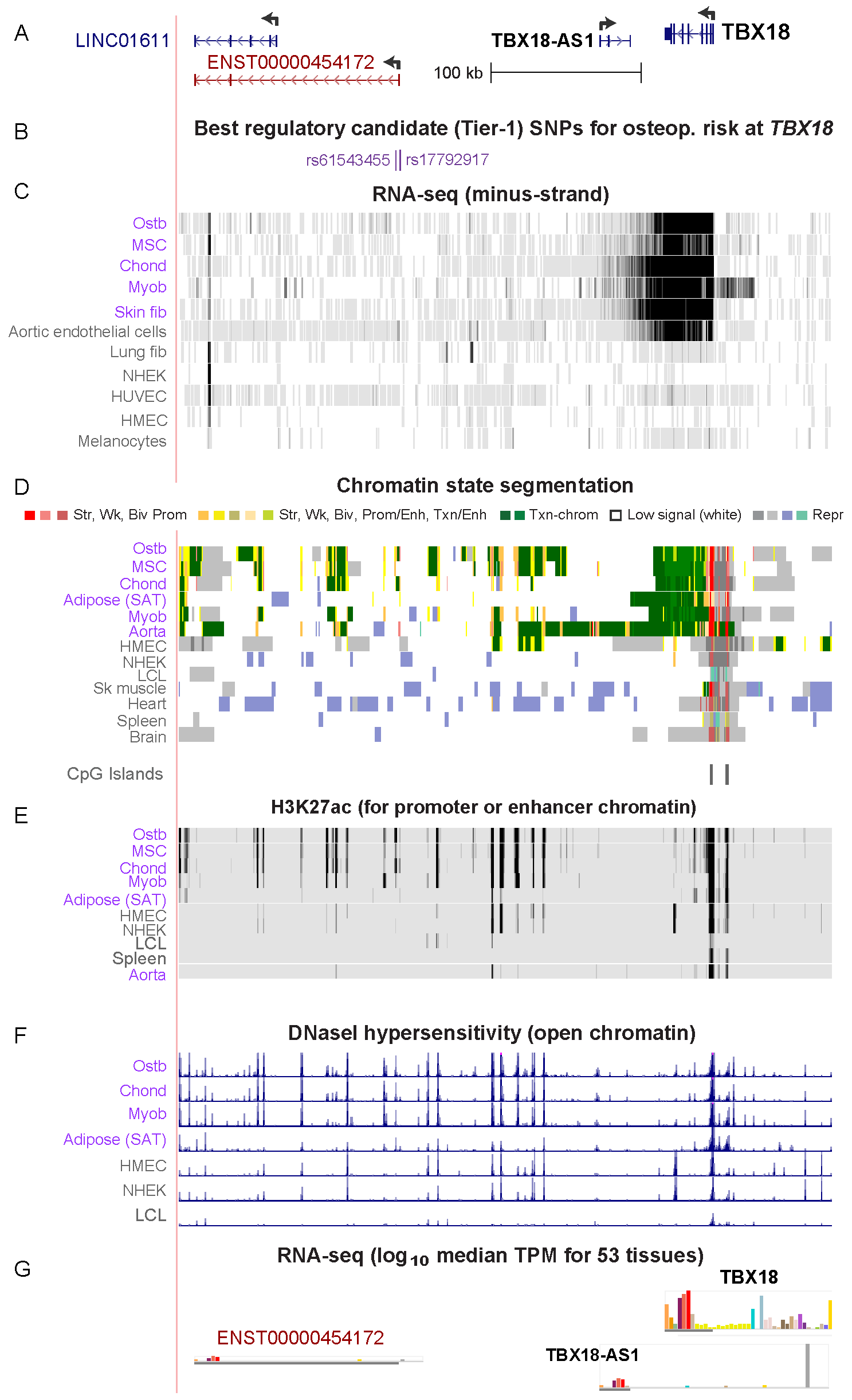
**Figure S3**. **Two eBMD Tier-1 SNPs are far downstream of *TBX18***. **(A)** RefSeq and ENSEMBL transcript genes in the *TBX18* neighborhood (chr6:85,120,602-85,552,957, hg19; UCSC Genome Browser). **(B)** Location of two eBMD Tier-1 SNPs. **(C)** Strand-specific RNA-seq for the indicated samples; vertical viewing scale 0-30. (**D-F)** Chromatin segmentation state, enrichment in H3K27 acetylation (vertical viewing range 0 – 4), and DNaseI hypersensitivity (vertical viewing range at 0 – 20) from the NIH Roadmap Epigenetics Project (5). **(G)** RNA-seq data from GTEx (6) showing similar tissue-specific expression for *TBX18* and the far downstream *ENST00000454172*. Abbreviations for tissues, cell types, and chromatin segmentation states are as described in Figs. 1-4 of the main article. Purple labels, *TBX18*-expressing cell populations; gray labels, not expressing.

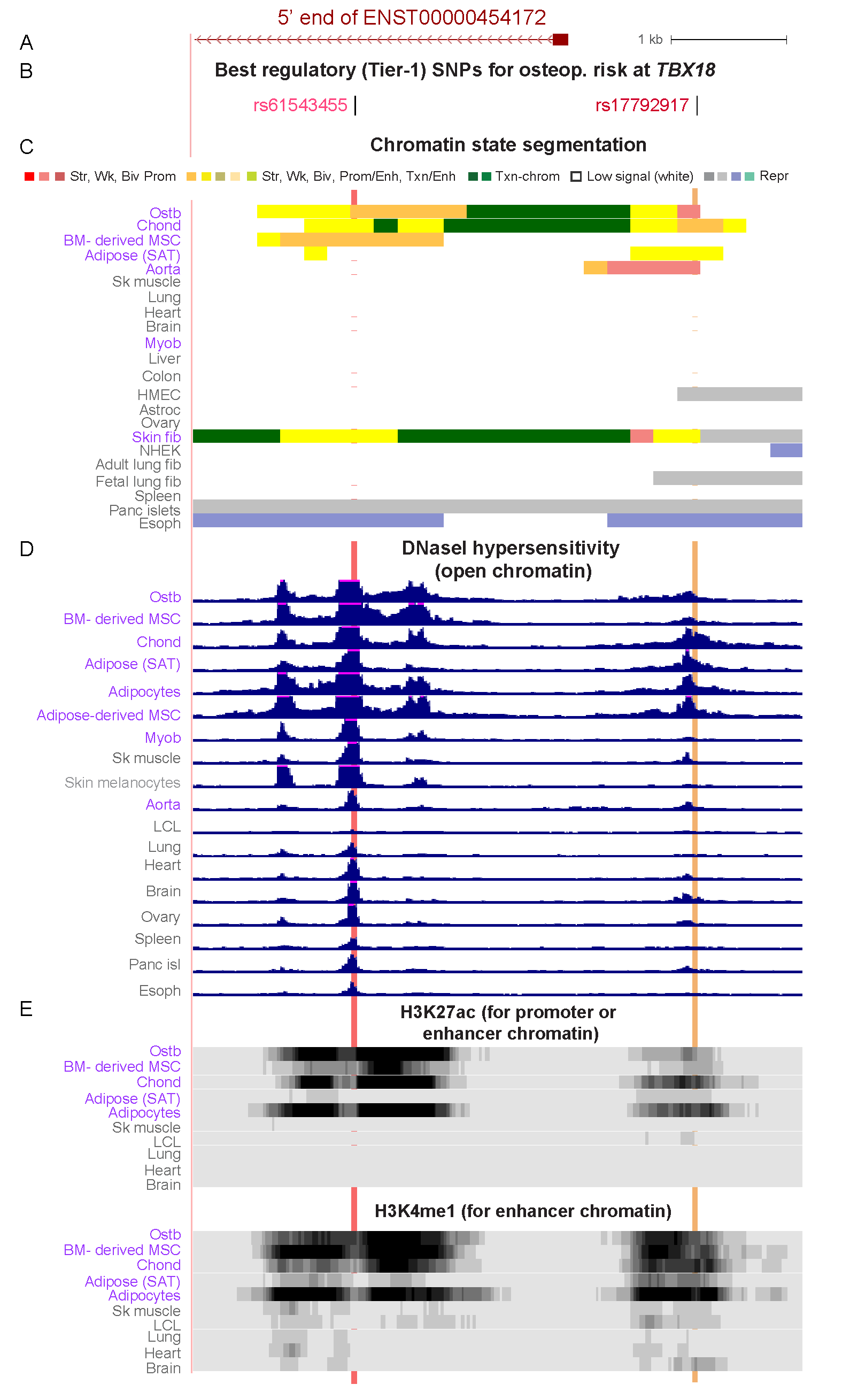
**Figure S4. Zoomed-in view of the two Tier-1 *TBX18* SNPs and their proximity to the 5’ end of a non-coding RNA gene preferentially transcribed in subcutaneous adipose (SAT) like *TBX18***. (**A – E)** Similar to panels in Fig. S3 except that the region shown is chr6:85,262,650-85,267,870 and the vertical viewing range for DNaseI hypersensitivity is 0 – 4. Highlighting, the positions of the two Tier-1 SNPs.
